# Proteomic profiling of *Mycobacterium tuberculosis* culture filtrate identifies novel O-glycosylated proteins

**DOI:** 10.1101/740134

**Authors:** Paula Tucci, Madelón Portela, Carlos Rivas Chetto, Gualberto González-Sapienza, Mónica Marín

## Abstract

Despite being the subject of intensive research, tuberculosis, caused by *Mycobacterium tuberculosis*, remains at present the leading cause of death from an infectious agent. Secreted and cell wall proteins interact with the host and play important roles in pathogenicity. These proteins have been explored as candidate diagnostic markers, potential drug targets or vaccine antigens, and special attention has been given to the role of their post-translational modifications. With the purpose of contributing to the proteomic characterization of this important pathogen including an O-glycosylation profile analysis, we performed a shotgun analysis of culture filtrate proteins of *M. tuberculosis* based on a liquid nano-HPLC tandem mass spectrometry and a label-free spectral counting normalization approach for protein quantification. We identified 1314 *M. tuberculosis* proteins in culture filtrate and found that the most abundant proteins belong to the extracellular region or cell wall compartment, and that the functional categories with higher protein abundance factor were virulence, detoxification and adaptation, and cell wall and cell processes. In culture filtrate, 140 proteins were predicted to contain one of the three types of bacterial N-terminal signal peptides. Besides, various proteins belonging to the ESX secretion systems, and to the PE and PPE families, secreted by the type VII secretion system using nonclassical secretion signals, were also identified. O-glycosylation was identified as a frequent modification, being present in 108 proteins, principally lipoproteins and secreted immunogenic antigens. We could identify a group of proteins consistently detected in previous studies, most of which were highly abundant proteins. Interestingly, we also provide proteomic evidence for 62 novel O-glycosylated proteins, aiding to the glycoproteomic characterization of relevant antigenic membrane and exported proteins.

## Introduction

*Mycobacterium tuberculosis*, the causative agent of tuberculosis (TB) remains a major public health threat. According to the last Global Tuberculosis Report published by the World Health Organization (WHO) an estimate of 10 million people developed TB disease in 2017. Moreover, TB is at present the leading cause of death from a single infectious agent, causing an estimated 1.3 million deaths among HIV-negative people and approximately 300 thousand deaths among HIV-positive people [1]. Although TB diagnosis and successful treatment averts millions of deaths each year, there are still large and persistent gaps related to this infection that must be resolved in order to accelerate progress towards the goal of ending the TB epidemic endorsed by WHO [1].

*M. tuberculosis* (MTB), has evolved successful mechanisms to circumvent the hostile environment of the macrophage, such as inhibiting the phagosome-lysosome fusion and to escape the acidic environment inside the phagolysosome [2]. MTB may be unique in its ability to exploit adaptive immune responses, through inflammatory lung tissue damage, to promote its transmission [3]. It has been proposed that this microorganism was pressed by an evolutionary selection that resulted in an infection that induces partial immunity, where the host survives a long period after being infected with the pathogen, aiding in microorganism persistence and transmission [3]. MTB mechanisms of evasion of host immune system were proposed to have consequences in the design of TB vaccines [3] and to be in part responsible of the poor performance of immune-based diagnostic tools [4,5].

In that context, there is a pressing need to advance the knowledge of the mechanisms that mediate its virulence. Among the tools to study the biology of MTB, *M. tuberculosis* H37Rv is a well-characterized human lung isolate and one of the most commonly used laboratory strains of *M. tuberculosis*. This virulent strain has been used in several investigations to understand the molecular mechanisms of MTB virulence, pathogenicity and persistence, as it provides a unique platform to investigate biochemical and signaling pathways associated with pathogenicity [6]. In particular, it has been extensively used to identify pathogen biomarkers of *M. tuberculosis* infection and disease. These are generally major components identified by electrophoresis and mass spectrometry both in total extracts and culture filtrates [7–9].

The cell envelope and secreted components of MTB are among the bacterial molecules most commonly described as potential biomarkers of the infection, or involved in host immune evasion. Mycobacteria possess a remarkably complex cell envelope consisting of a cytoplasmic membrane and a cell wall. These constitute an efficient permeability barrier that plays a crucial role in intrinsic drug resistance and contributes to the resilience of the pathogen in infected hosts [10]. Membrane and exported proteins are crucial players for maintenance and survival of bacterial organisms, and their contribution to pathogenesis and immunological responses make these proteins relevant targets for medical research [11]. In particular, these proteins are known to play pivotal roles in host-pathogen interactions and, therefore, represent potential drug targets and vaccine candidates [12].

Overall, the bulk of exported proteins are transported by the general secretory Sec-translocase pathway. This is performed by recognition of the signal peptide in the nascent preprotein, which is subsequently transferred to the machinery that executes its translocation across the membrane [13]. Besides, mycobacteria utilize type VII secretion systems (T7SS) to export many of their important virulence proteins. The T7SS encompasses five homologous secretion systems (designated ESX-1 through ESX-5). Most pathogenic mycobacterial species, including the human pathogen *M. tuberculosis*, possess all five ESX systems [14,15]. The ability of MTB to subvert host immune defenses is related to the secretion of multiple virulence factors via these specialized secretion systems [15].

Recent developments in mass spectrometry-based proteomics have highlighted the occurrence of numerous types of post-translational modifications (PTMs) in proteomes of prokaryotes which create an enormous diversity and complexity of gene products [16]. This PTMs, mainly glycosylation, lipidation and phosphorylation, are involved in signaling and response to stress, adaptation to changing environments, regulation of toxic and damaged proteins, protein localization and host-pathogen interactions. In MTB, more frequently O-glycosylation events have been reported [17], being this post-translational modification often found, in conjunction with acylation, in membrane lipoproteins [18]. A mechanistic model of this modification was proposed in which the initial glycosyl molecule is transferred to the hydroxyl oxygen of the acceptor Thr or Ser residue, a process catalyzed by the protein O-mannosyltransferase (PMT) (Rv1002c) [19]. Hereafter, further sugars are added one at a time, but the enzymes involved in this elongation are still unknown [16]. O-glycosylation appears essential for MTB virulence, since Rv1002c deficient strains are highly attenuated in immunocompromised mice [20]. Despite the vital importance of glycosylated proteins in MTB pathogenesis, the current knowledge in this regard is still limited. Recent evidence using whole cell extracts revealed that glycosylation could be much more frequent than previously thought, explaining the phenotypic diversity and virulence in the *Mycobacterium tuberculosis* complex [17], but in culture filtrates of this pathogen only a few secreted and cell wall-associated glycoproteins have been described to date [18,21].

In this study we describe a straightforward methodology based on a high throughput label-free quantitative proteomic approach in order to provide a comprehensive identification and quantitation of proteins in *M. tuberculosis* H37Rv culture filtrate. The extent of protein O-glycosylation was also evaluated with the purpose of collaborating with the glycoproteomic characterization of this pathogen. With the goal to validate and integrate our results, a comprehensive comparative analysis was performed against former research papers that have addressed this issue using different and complementary approaches. The results presented here make focus on the principal exported and secreted virulent factors with the aim to contribute to a deep proteomic characterization of this relevant pathogen and to collaborate to a better understanding of the pathogenesis and survival strategies adopted by MTB.

## Materials and Methods

### Mycobacterial strain and growth conditions

*Mycobacterium tuberculosis* H37Rv strain (ATCC® 25618™) was grown for 3 weeks at 37°C in Lowenstein Jensen solid medium and after growth was achieved it was subcultured in Middlebrook 7H9 broth supplemented with albumin, dextrose, and catalase (ADC) enrichment (Difco, Detroit, MI, USA) for 12 days with gentle agitation at 37°C. Mycobacterial cells were pelleted at 4000xg for 15 min at 4°C and washed 3 times with cold phosphate-buffered saline. Mycobacterial cells were subsequently cultured as surface pellicles for 3 to 4 weeks at 37°C without shaking in 250 mL of Sauton minimal medium, a synthetic protein-free culture medium, which was prepared as previously described [8].

### Culture filtrate protein preparation

Bacterial cells were removed by centrifugation and culture filtrate protein (CFP) was prepared by filtering the supernatant through 0.2 µM pore size filters (Millipore, USA). After sterility testing of CFP in Mycobacteria Growth Indicator Tube (MGIT) supplemented with MGIT 960 supplement (BD, Bactec) for 42 days at 37°C in BD BACTEC™ MGIT™ automated mycobacterial detection system, CFP was concentrated using centrifugal filter devices (Macrosep Advance, 3kDa MWCO (Pall Corporation, USA)). Concentrated CFP was buffer exchanged to phosphate-buffered saline and total protein concentration was quantified by BCA (Pierce BCA Protein Assay Kit, Thermo Fischer Scientific).

### 1D and 2D gel electrophoresis

*M. tuberculosis* CFP samples were analyzed by 1D and 2D gel electrophoresis and were used for raising polyclonal antibodies in rabbits as described below. For 1D gel electrophoresis CFP diluted in SDS-PAGE loading buffer was loaded onto 15% SDS-PAGE and silver nitrate staining was performed as described elsewhere [22]. For 2-Dimensional gel electrophoresis 50 µg of *M. tuberculosis* CFP was purified and concentrated using 2-D Clean-Up Kit (GE Healthcare) and resuspended in 125 µl of rehydration solution (urea 7M, thiourea 2M, CHAPS 2%, IPG Buffer 3-10 0,5%, DTT 20 mM, bromophenol blue 0,002%). Two experiments were run in parallel, one for silver nitrate staining and the other for western blot analysis. Proteins were loaded into 7 cm IPG Strips 3-10 (GE Healthcare) by overnight passive rehydration. First dimension isoelectric focusing (IEF) run was performed using Ettan IPGphor 3 IEF System (GE Healthcare) according to manufacturer instructions. Disulfide bonds were reduced with dithiothreitol (10 mg/mL) and subsequently alkylated with 25 mg/mL iodoacetamide. The second dimension was performed on hand-cast gels (15% SDS-PAGE, 10×10×0.1cm) and silver nitrate staining was performed as described above. In both analyses the molecular weight marker was PageRuler Prestained Protein Ladder (Thermo Fischer Scientific). For Western blot analysis, proteins were transferred onto a nitrocellulose membrane (Amersham Protran 0.45 µM NC (GE Healthcare)) for one hour at 400mA. Membrane was blocked and blotted as described below.

### Anti-CFP antibodies production and western blot

To produce polyclonal antibodies against *M. tuberculosis* CFP, two New Zealand White rabbits (2-2.5 kg) were immunized subcutaneously with 100 µg of CFP, followed by 1 booster of 100 µg and 2 additional boosters (50 µg each) of CFP in Incomplete Freund Adjuvant using an authorized protocol (Comité de Etica de Facultad de Química, Exp. № 101900-000717-14). At the end the rabbits were bled and a pool of hyperimmune serum was obtained as described elsewhere [23]. Anti-CFP polyclonal antibodies were purified by affinity chromatography using a HiTrap Protein A HP column (GE Healthcare) according to manufacturer instructions.

Anti-CFP polyclonal antibodies were used to identify immunoreactive bands and spots in 1D and 2D electrophoresis. Briefly, blocked membranes were incubated for 1h at room temperature with anti-CFP antibodies at a final concentration of 10 μg/mL in PBS pH7.4, 5% low fat milk. For antigen-antibody detection membranes were incubated for 1h at room temperature with a 1:2500 dilution of anti-rabbit IgG (whole molecule)— alkaline phosphatase antibody produced in goat (Sigma A0545) in PBS pH7.4, 5% low fat milk. Membranes were incubated with SuperSignal™ West Pico PLUS Chemiluminescent Substrate (Thermo Fischer Scientific, #34580) according to manufacturer instructions. Images were acquired with Synoptics 4.2MP Camara using increasing and accumulative exposure times in G:Box Chemi XT4 (Syngene, Cambridge, UK) and visualized with GenSys Software (V1.3.3.0). The following reagent, obtained through BEI Resources, NIAID, NIH: Polyclonal Anti-Mycobacterium tuberculosis CFP minus LAM (antiserum, Rabbit), NR-13809, was used to confirm the immune recognition of our Anti-CFP polyclonal antibody. Some of the protein spots recognized by the anti-CFP antibody were further analyzed by mass spectrometry (MS).

### Protein identification by MALDI-TOF/TOF

Bands or spots from 1D or 2D gel electrophoresis were selected for MS MALDI-TOF/TOF analysis. In-gel Cys alkylation was performed by subsequent incubation with 10 mM dithiothreitol and 55 mM iodoacetamide as previously described [24]. In-gel digestion of selected protein bands or spots was performed overnight at 37 °C by incubation with trypsin (Sequencing grade, Promega, Madison, USA). Afterwards peptides were extracted as previously described [24] and samples were vacuum-dried using CentriVap Vacuum Concentrator (Labconco), resuspended in 0.1% TFA, and desalted using C18 OMIX tips (Agilent). Peptides were eluted with matrix solution (α-cyano-4-hydroxycinnamic acid in 60% acetonitrile, 0.1% TFA) directly into the MALDI sample plate. Spectra acquisition was performed on a 4800 MALDI TOF/TOF (Abi Sciex) operating in positive reflector mode. Spectra were externally calibrated using a mixture of peptide standards (Applied Biosystems).

MS/MS analysis of selected precursor ions was performed. Database searching (NCBInr 20150912) was performed with Mascot (http://www.matrixscience.com) using the following parameters: unrestricted taxonomy; one trypsin missed cleavage allowed; methionine oxidation and carbamidomethylation of cysteine as variable modification; peptide tolerance of 0.05 Da and a MS/MS tolerance of 0.4 Da. Significant protein scores (p <0.05) and at least one peptide with significant ions score (p <0.05) per protein were used as criteria for positive identification [25].

### Liquid chromatography tandem mass spectrometry (LC MS/MS)

Two replicas of *M. tuberculosis* CFP (25 µg) were loaded in SDS-PAGE 15% and stained with CCB G-250 as described elsewhere [26]. Six gel slices were excised from each lane according to protein density. In-gel Cys alkylation, in gel-digestion and peptide extraction was performed as described above. Tryptic peptides were separated using nano-HPLC (UltiMate 3000, Thermo Scientific) coupled online with a Q-Exactive Plus hybrid quadrupole-Orbitrap mass spectrometer (Thermo Fischer Scientific). Peptide mixtures were injected into a trap column Acclaim PepMap 100, C18, 75 um ID, 20 mm length, 3 um particle size (Thermo Scientific) and separated into a Reprosil-Pur 120 C18-AQ, 3 µm (Dr. Maisch) self-packed column (75µm ID, 49 cm length) at a flow rate of 250 nL/min. Peptide elution was achieved with 105 min gradient from 5% to 55% of mobile phase B (A: 0.1% formic acid; B: 0.1% formic acid in 80% acetonitrile). The mass spectrometer was operated in data-dependent acquisition mode with automatic switching between MS and MS/MS scans. The full MS scans were acquired at 70K resolution with automatic gain control (AGC) target of 1 × 10^6^ ions between m/z = 200 to 2000 and were surveyed for a maximum injection time of 100 milliseconds (ms). Higher-energy collision dissociation (HCD) was used for peptide fragmentation at normalized collision energy set to 30. The MS/MS scans were performed using a data-dependent top12 method at a resolution of 17.5K with an AGC of 1 × 10^5^ ions at a maximum injection time of 50 ms and isolation window of 2.0 m/z units. A dynamic exclusion list with a dynamic exclusion duration of 45 s was applied.

### LC-MS/MS data analysis

LC-MS/MS data analysis was performed in accordance to the PatternLab for proteomics 4.0 software (http://www.patternlabforproteomics.org) data analysis protocol [27].The proteome (n=3993 proteins) from *M. tuberculosis* (Reference strain ATCC 25618/H37Rv UP000001584) was downloaded from Uniprot (March 2017) (https://www.uniprot.org/proteomes/). A target-reverse data-base including the 123 most common contaminants was generated using PatternLab’s database generation tool. Thermo raw files were searched against the database using the integrated Comet [28] search engine (2016.01rev.3) with the following parameters: mass tolerance from the measured precursor m/z(ppm): 40; enzyme: trypsin, enzyme specificity: semi-specific, missed cleavages: 2; variable modifications: methionine oxidation; fixed modifications: carbamidomethylation of cysteine. Peptide spectrum matches were then filtered using PatternLab’s Search Engine Processor (SEPro) module to achieve a list of identifications with less than 1% of false discovery rate (FDR) at the protein level [29]. Results were post-processed to only accept peptides with six or more residues and proteins with at least two different peptide spectrum matches. These last filters led to an FDR at the protein level, to be lower than 1% for all search results. Proteins were further grouped according to a maximum parsimony criteria in order to identify protein clusters with shared peptides and to derive the minimal list of proteins [30]. Spectrum counts of proteins identified in each technical replicate were statistically compared with unpaired Mann-Whitney test.

For the O-glycosylation analysis raw files were searched against the same database using the parameters described above with the addition of the following variable modifications in S or T amino acid residues: Hex =162.052824 Da, Hex-Hex=324.1056 Da, Hex-Hex-Hex=486.1584 Da, Pentose=132.042259 Da, Heptose=192.0633 Da, DeoxyHex=146.0579 Da. Monoisotopic mass of each neutral loss modification was defined in Comet search engine according to the values recorded in Unimod public domain database (http://www.unimod.org/). Each O-glycosylation was tested independently and a maximum of 2 modifications per peptide was allowed.

Peptide spectrum matches were filtered and post-processed using SEPro module, using the same parameters as described above and proteins were grouped according to a maximum parsimony criteria [30].

### Protein analysis

Identified proteins in each replicate were compared by area-proportional Venn Diagram comparison (BioVenn [31]) and a list of common proteins was generated. Further analysis only considered proteins present in both replicates of LC MS/MS analysis. SEPro module retrieved a list of protein identified with Uniprot code. Molecular weight, length, complete sequence, gene name and *M. tuberculosis* locus identified (Rv) was obtained using the Retrieve/ID mapping Tool of Uniprot website (https://www.uniprot.org/uploadlists/) [32]. Protein functional category was obtained by downloading *M. tuberculosis* H37Rv genome sequence Release 3 (2018-06-05) from Mycobrowser website (https://mycobrowser.epfl.ch/) [33].

### Protein O-glycosylation analysis

Proteins bearing O-glycosylated peptides in both replicates were compared by area-proportional Venn Diagram comparison (BioVenn [31]) and a list of common glycosylated proteins for each of the analyzed modifications, i.e. Hex, Hex-Hex, Hex-Hex-Hex, Pentose, Heptose, DeoxyHex, was generated. Further analysis was manually performed in order to identify common modified peptides in the list of common glycosylated proteins, as well as common modifications (as 1 peptide could contain up to two modifications). As a result of this analysis a list of proteins with common modifications was generated, consisting in proteins having the same modified peptide in both replicates. This list of O-glycosylated proteins was considered for subsequent analysis.

### Signal peptide prediction

In order to identify potentially secreted proteins, the SignalP 5.0 Server (http://www.cbs.dtu.dk/services/SignalP/) was used to detect the presence of N-terminal signal sequences in the analyzed set of proteins. The organism group selected was gram-positive bacteria. This version of the Server, recently launched, can predict proteome-wide signal peptides across all organisms, and classify them into three type of signal peptides: Sec/SPI (SP), Sec/SPII (LIPO) and Tat/SPI (TAT) [34]. In the output produced by the server one annotation is attributed to each protein, the one that has the highest probability. The protein can have a Sec signal peptide (Sec/SPI), a Lipoprotein signal peptide (Sec/SPII), a Tat signal peptide (Tat/SPI) or No signal peptide at all (Other). If a signal peptide is predicted, the cleavage site (CS) position is also reported.

### Estimation of protein abundance and comparative analysis

To estimate protein abundance Normalized Spectral Abundance Factor (NSAF) calculated with PatternLab for proteomics software was considered. NSAF allows for the estimation of protein abundance by dividing the sum of spectral counts for each identified protein by its length, thus determining the spectral abundance factor (SAF), and normalizing this value against the sum of the total protein SAFs in the sample [35,36]. Proteins were ordered according to their NSAF, from more to less abundant. NSAF values corresponding to percentile 75th, 90th and 95th were calculated, and the groups of proteins above these values were identified as P75%, P90% and P95% proteins, respectively. The list of proteins obtained in this study was compared with other proteomic studies [7,13,37] by Venn Diagram comparison (Venny 2.1, BioinfoGP [38]) and NSAF of proteins identified in all studies, 3 studies, 2 studies or only this study were statistically compared with unpaired Mann-Whitney test. The protein abundance determined for CFP identified in this study (NSAF) was compared with the protein abundance calculated for *M. tuberculosis* proteins identified in a previous study using the exponentially modified protein abundance index (emPAI) [13].

### Protein classification

Gene Onthology (GO) analysis of the culture filtrate proteins was performed with David Gene Functional Classification Tool [39,40] using the Cellular Component Ontology database and *M. tuberculosis* H37Rv total proteins as background. With this analysis principal categories of enriched terms (p<0.05) for P75%, P90%, P95% and total proteins were determined. Functional classification of culture filtrate proteins was performed according to functional categories of *M. tuberculosis* knowledge database (Mycobrowser [33]).

Proteins with O-glycosylation modifications were analyzed with David Gene Functional Classification Tool [39,40] using Cellular Component, Biological Processes and Molecular functions Ontology database and *M. tuberculosis* H37Rv total proteins as background.

### O-glycosylation validation

The same analytical workflow described previously for LC-MS/MS analysis of O-glycosylation in our data was performed using the raw data files deposited at the ProteomeXchange Consortium with the dataset identifier PXD000111 [37]. This analysis was performed in order to validate the modified peptides identified in our work against additional biological replicates obtained in a previous work that extensively characterized culture filtrate proteins of *M. tuberculosis* H37Rv [37]. Additionally, some relevant scans corresponding to glycosylated peptides were searched in Mascot Server MS/MS Ions Search (Mascot, Matrix Science Limited [41]). Search was performed against NCBIprot (AA) database of all taxonomies. Search parameters were defined as peptide mass tolerance: ± 10 ppm, MS/MS mass tolerance: ± 0.15 Da, enzyme: semiTrypsin, fixed modifications: Carbamidomethyl (C), variable modifications: Hex (ST), Hex(2) (ST), Hex(3) (ST), Pent (ST), Hept (ST) or dHex (ST), according to the searched peptide. Other parameters were set to default values.

## Results and Discussion

### *M. tuberculosis* culture filtrate proteins quality evaluation

*M. tuberculosis* H37Rv was cultured following a classical method using Sauton minimal medium, a synthetic protein-free culture medium compatible with proteomic downstream analysis [8]. Culture filtrate proteins (CFP), obtained after culture centrifugation and filtration, were concentrated by ultrafiltration and quantitated previous to further analysis. Four different batches of CPF were analyzed by gel electrophoresis and silver nitrate staining. As similar patterns were observed with the different CFP preparations a composed sample was prepared. The composed CFP sample was separated and resolved by 1D and 2D gel electrophoresis. In 1D SDS-PAGE an electrophoretic pattern showing a variety of proteins from approx. 10 kDa to 100 kDa was observed (Fig 1A). In the case of 2D gel electrophoresis analysis, two experiments were run in parallel, one for silver nitrate staining (Fig 1B) and the other for western blot analysis using anti-CFP rabbit polyclonal antibodies to identify principal immunogenic proteins (Fig 1C). Immune recognition pattern of CFP proteins observed with anti-CFP rabbit polyclonal antibody was confirmed with an additional anti-CFP polyclonal antibody (NR-13809, BEI Resources).

**Fig 1.**
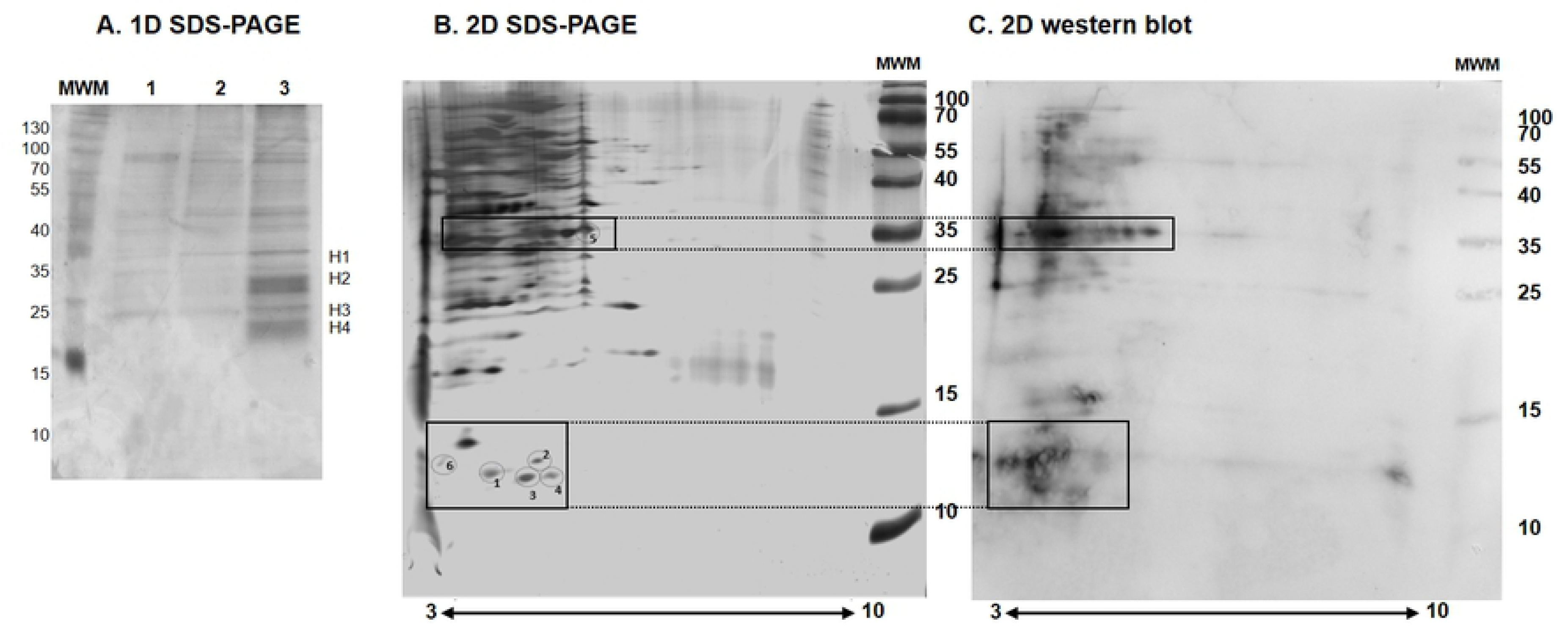
Analysis of *M. tuberculosis* CFP by electrophoresis and *western blot*. (A) *M. tuberculosis* CFP analysis by 1D SDS-PAGE 15% and silver nitrate staining. Two different batches (lanes 1 and 2, 1.8 and 2.1 ug, respectively) and a composed and concentrated sample of both batches (lane 3, 12 ug) were loaded. Bands selected for MALDI-TOF/TOF mass spectrometry are indicated as H1, H2, H3 and H4. MWM: Molecular weight marker (Thermo Fischer Scientific, # 26616). (B) *M. tuberculosis* CFP analysis by 2D electrophoresis and silver nitrate staining. *M. tuberculosis* CFP composed sample (50 ug) was loaded. Immunoreactive spots selected for MALDI-TOF/TOF mass spectrometry are indicated with numbers. MWM: Molecular weight marker (Thermo Fischer Scientific, # 26616). (C) Western blot analysis of *M. tuberculosis* CPF. 2D gel performed equally as (B) was transferred to Protran 0.45 uM NC (GE Healtcare) and probed with rabbit anti-CFP antibody. Immunoreactive zones are indicated with a rectangle in the corresponding 2D gel.

As shown in Fig 1B most of the spots consisted of proteins with an isoelectric point below 6.5, as was previously reported by others [7,8,42,43]. Some immunoreactive spots detected in 2D western blot were overlapped with 2D gel silver nitrate stained to select candidates to be analyzed by mass spectrometry (Fig 1C). By this MS analysis 12 different proteins of *M. tuberculosis* (MTB) were identified in the CFP sample (Table 1) as well as some low-signal contaminant keratin peptides. Molecular weight of identified MTB proteins showed a good correlation with the relative molecular weight of selected band or spot (Table 1).

**Table 1.** M. tuberculosis proteins identified by MALDI-TOF (MS/MS) from 1D and 2D SDS polyacrylamide electrophoresis.

| Band / spot | Protein | Molecular weight | Gene name | Gene identifier | Proteomics | Functional category |
| --- | --- | --- | --- | --- | --- | --- |
| **H1** | Conserved protein with FHA domain, GarA | 17,2 kDa | *garA* | **Rv1827** | <u>CF</u>, CYT, CW, MF. | Conserved hypothetical |
| **H1** | Adenylate kinase Adk (ATP-AMP transphosphorylase) | 20,0 kDa | *adk* | **Rv0733** | <u>CF</u>, CYT, MF. | Intermediary metabolism and respiration |
| **H1** | Superoxide dismutase [FE] SodA | 23,0 kDa | *sodA* | **Rv3846** | <u>CF</u>, CYT, MF. | Virulence, detoxification, adaptation |
| **H2** | Conserved protein TB18.6 | 18,6 kDa | TB18.6 | **Rv2140c** | <u>CF</u>, MF. | Conserved hypotheticals |
| **H2** | Probable thiol peroxidase Tpx | 16,8 kDa | *tpx* | **Rv1932** | <u>CF</u>, CYT, MF. | Virulence, detoxification, adaptation |
| **H3** | 10 kDa chaperonin GroES | 10,8 kDa | *groES* | **Rv3418c** | <u>CF</u>, CYT, CW, MF. | Virulence, detoxification, adaptation |
| **H3/1/2/6** | Heat shock protein HspX (alpha-crystallin homolog) | 16,2 kDa | *hspX* | **Rv2031c** | <u>CF</u>, CYT, CW, MF. | Virulence, detoxification, adaptation |
| **H4/1/6** | 10 kDa culture filtrate antigen EsxB | 10,8 kDa | *esxB* | **Rv3874** | <u>CF</u>, CYT, MF. | Cell wall and cell processes |
| **1/2/3/4** | ESAT-6 like protein (Identification could correspond to EsxJ, EsxK, EsxM, EsxP or EsxW) | ≅ 10 kDa | *esxJ*<br>*esxK*<br>*esxM*<br>*esxP*<br>*esxW* | **Rv1038c**<br>**Rv1197**<br>**Rv1792**<br>**Rv2347c**<br>**Rv3620c** | <u>CF</u>* | Cell wall and cell processes |
| **2** | Thioredoxin TrxC (TRX) (MPT46) | 12.5 kDa | *trxC* | **Rv3914** | <u>CF</u>, CYT, CW, MF. | Intermediary metabolism and respiration |
| **5** | Secreted antigen 85-a FbpA (mycolyl transferase 85A) | 35.7 kDa | *fbpA* | **Rv3804c** | <u>CF</u>, CYT, CW, MF. | Lipid metabolism |
| **6** | Rv3747 | 13.5 kDa | Rv3747 | **Rv3747** | MF | Conserved hypotheticals |

CF: Culture filtrate, CYT: Cytosol, CW: Cell Wall, MF: Membrane fraction, \* Putative (EsxK, EsxM), reported (EsxJ, EsxW), not reported (EsxP). Table filled with information obtained from Ref [33].

All proteins identified by this approach were previously detected in other proteomic studies, and most of them (11 out of 12) were identified in the culture filtrate fraction by at least one earlier proteomic report [33]. Besides, 3 proteins were identified in at least 3 different bands or spots, reflecting the fact that these proteins could be highly represented in CFP. These are heat shock protein HspX (Rv2031c), 10 kDa culture filtrate antigen EsxB (Rv3874) and a group of indistinguishable proteins (ESAT-6 like proteins). As proteins of this group - EsxJ (Rv1038c), EsxK (Rv1197), EsxM (Rv1792), EsxP (Rv2347c) and EsxW (Rv3620c) - are 98 amino acids long and differ in only 1 or two amino acids, the unequivocal identification of each one is hindered. Proteins of ESAT-6 like group were identified in a common zone of the 2D gel (spots 1 to 4), suggesting that each spot could correspond to a slightly different protein isoform.

These results indicated that the *M. tuberculosis* H37Rv CFP preparation provides a good representation of the secreted/shed proteins because many main proteins of MTB were identified, with minimal contamination of non-MTB proteins (only a few peptides of human keratin were detected). Proteins highly recognized by anti-CFP antibodies (HspX, EsxB and Secreted antigen 85-a FbpA (Rv3804c)) are relevant pathogen antigens [33,44], recognized as secreted proteins by others [8,13], and evaluated as pathogen-derived biomarkers for active tuberculosis diagnosis [9].

### Characterization of CFP using LC MS/MS

Although 2D gel electrophoresis coupled with MALDI TOF/TOF analysis is an extremely powerful tool to dissect and resolve multiprotein complexes, it is a low performance methodology for proteomic analysis, which needs a laborious and systematic approach in order to get confident and sensitive identification of proteins present in complex samples. Our results showed that this analysis generated several cases of redundant identifications. Thus, after quality confirmation of the sample, a high throughput analysis was performed using a shotgun quantitative approach based on a liquid nano-HPLC and tandem mass spectrometry workflow. In this experiment the proteins present in two technical replicates were resolved in SDS-PAGE and different portions of the gel were further selected for LC MS/MS analysis (S1A Fig). A gross initial quantitative comparison of spectrum counts of both datasets showed that there were not statistical differences among both replicates (S1B Fig). In CFP(1) 1450 different proteins were identified (corresponding 1427 to MTB, 19 to common contaminants and 4 to reverse sequences, resulting in a 0.28% FDR), whereas in CFP(2) 1453 different proteins were identified (1429 MTB proteins, 18 contaminants and 6 reverse sequences (0.41% FDR)). The list of proteins of each replica is available in S1 Table. The mass spectrometry proteomics data (raw data and search files) have been deposited at the MassIVE repository with the dataset DOI: doi:10.25345/C5PW8Q.

The qualitative comparison of both datasets using a Venn Diagram bioinformatic tool showed that 1314 MTB proteins (92%) were shared between both replicates (S1C Fig). All proteins previously identified in the CFP sample by gel electrophoresis and MALDI-TOF/TOF were detected in both replicates characterized by LC MS/MS. The full list of 1314 common proteins, which was used for further analysis, is provided in S1 Table. Proteins showed a wide distribution of molecular weights, however most of them were of low molecular weight (median 31.97 kDa, Q1 21.25 kDa, Q3 46.50 kDa), which was consistent with the profile observed in Fig 1A and 1B. Previous research has shown that the vast majority of protein spots resolved in 2D gel electrophoresis of *M. tuberculosis* H37Rv CFP were found in the molecular weight range of 6–70 kDa [8]. Moreover, consistent with our results, proteins identified by LC-MS/MS in a well characterized CFP, showed that the majority of the proteins were found in the 10-50 kDa range, with an average theoretical mass of 31.0 kDa [7].

### Protein classification using a quali-quantitative analysis

Quantitative proteomics based on spectral counting methods are straightforward to employ and have been shown to correctly detect differences between samples [45]. In order to consider sample-to-sample variation obtained when carrying out replicate analyses, and due to the fact that longer proteins tend to have more peptide identifications than shorter proteins, Patternlab for Proteomics software uses NSAF (Normalized spectral abundance factor) [46] for spectral counting normalization. The NSAF for a protein is the number of spectral counts (SpC, the total number of MS/MS spectra) identifying a protein, divided by the protein’s length (L), divided by the sum of SpC/L for all N proteins in the experiment. NSAF was shown to yield the most reproducible counts across technical and biological replicates [35]. Using the sum of NSAF of both replicates (Total NSAF, included in S1 Table) the common list of CFP was ordered according to protein abundance and arbitrarily grouped in 4 subgroups (P95%, P90%, P75% and total CFP), consisting of 66, 132, 329 and 1314 proteins, respectively. P95% comprised proteins above 95th percentile NSAF, thus representing the most abundant proteins in the sample. P90% and P75% comprised proteins above 90th and 75th percentile, respectively. These subgroups of proteins were functionally classified using Gene Ontology, Cellular Component analysis, and principal categories of enriched terms (p<0.05) were determined (Fig 2A). Considering the subgroup of total CFP proteins 4 principal categories (cell wall, cytoplasm, extracellular region and plasma membrane) were similarly enriched (fold change 1.5, 1.5, 1.2 and 1.1, respectively). However, when considering the subgroups of more abundant proteins, the categories cell wall and extracellular region showed a marked increase of fold enrichment with protein abundance, achieving these categories in P95% subgroup a fold enrichment of 2.9 (p=8.3e-18) and 3.1 (p=2.0e-8), respectively. This tendency was not observed in cytoplasm and plasma membrane categories.

**Figure 2.**
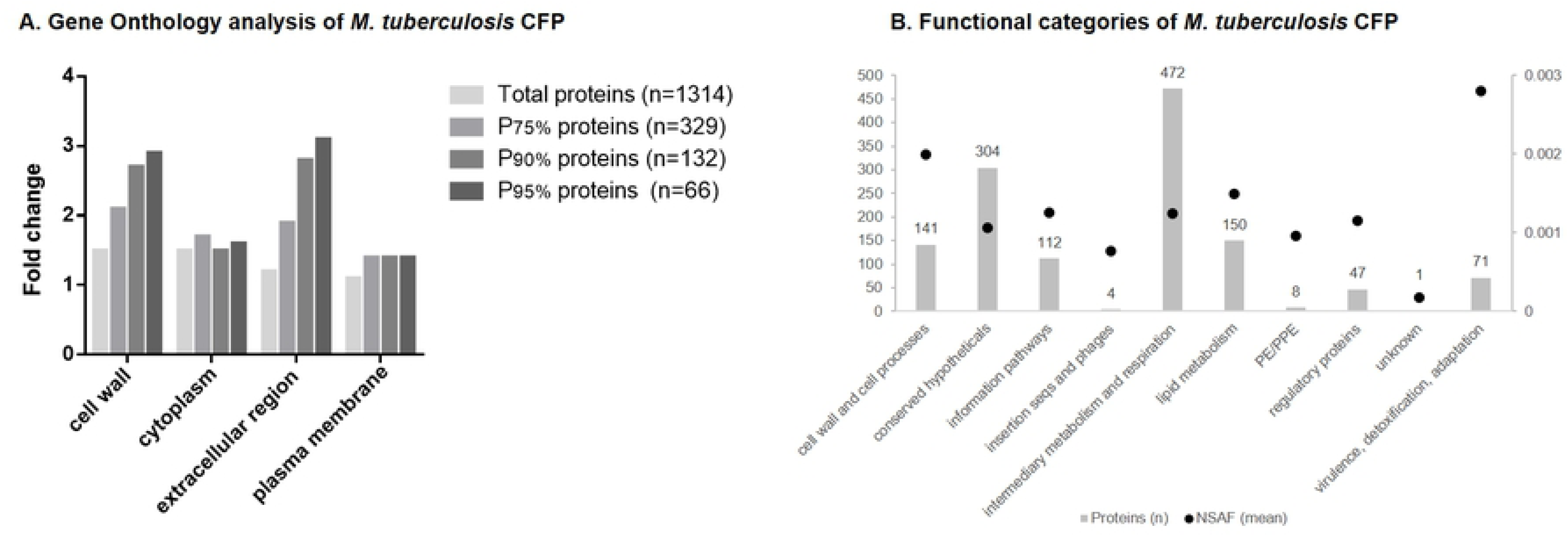
Quali-quantitative protein classification. (A) Fold change of principal categories of enriched terms (p<0.05) obtained analyzing common proteins of both replicates with David Gene Functional Classification Tool [39,40] using the Cellular Component Ontology database and *M. tuberculosis* H37Rv total proteins as background. Proteins were ordered considering normalized spectral abundance factor (NSAF) and percentile 75th, 90th and 95th NSAF were calculated. Fold change of the lists above each defined percentile (P75%, P90% and P95% proteins) analyzed using the same approach is shown. (B) Functional categories of CFP according to *M. tuberculosis* knowledge database (Mycobrowser [33]). Bars represent number of proteins corresponding to each category (number is indicated above each bar, scale in left axe) and dots represent mean NSAF of proteins in each category (scale is indicated in right axe).

The results presented showed that CFP proteins prepared in this work besides containing extracellular and cell wall proteins also include some cytoplasmatic and membrane proteins. This observation should be relativized considering the fact that many CFP were classified with more than one ontology term, thus redundant information of cellular component could be obtained. Particularly, 183 proteins were classified as extracellular, but only 44 contained exclusively this ontology term. Besides, only 125 proteins were classified as exclusively cytoplasmatic, out of 463 proteins containing this ontology term. It is also important to note that 394 proteins had no assigned GO term. Taken this into account the analysis performed considering the abundance of each CFP protein in terms of NSAF could be more indicative of the actual composition of the sample. In that regard, our analysis indicates that the subgroups of more abundant proteins contained mainly proteins of extracellular region and cell wall compartment.

The annotated *M. tuberculosis* H37Rv proteins have been classified into 12 distinct functional categories in the *M. tuberculosis* knowledge database (Mycobrowser [33]). Functional classification of proteins identified in this study according to this classification showed that proteins were distributed across ten of those functional groups (Fig 2B). Most of the identified proteins are involved in intermediary metabolism and respiration (35.9%). However, when protein abundance is considered, the category with higher protein mean NSAF is virulence, detoxification, adaptation followed by cell wall and cell processes (Fig 2B).

Finally, considering the need of pathogen-derived biomarker validation for *M. tuberculosis* active diagnosis, we looked in the list of CFP for principal protein antigens detected in clinical samples [9], confirming the presence of 11 out of 12. Moreover, these putative biomarkers exhibited on average a high NSAF, being 10 of them in the P90% subgroup: GroEL2 (Rv0440), EsxA (Rv3875), HspX (Rv2031c), FbpA (Rv3804c), FbpB (Rv1886c), Mpt64 (Rv1980c), PstS1 (Rv0934), GlcB (Rv1837c), Apa (Rv1860) and FbpC (Rv0129c).

### Prediction of secreted proteins

Given the results obtained the question arises whether the presence of certain proteins in CFP is due to bacterial leakage/autolysis in combination with high levels of protein expression and extracellular stability, rather than to protein-specific export mechanisms. *M. tuberculosis* H37Rv reference proteome (UP000001584) obtained from UniProt and our list of proteins from culture filtrate was submitted to SignalP 5.0 signal peptide prediction [34]. This method incorporates deep recurrent neural network-based approach that improves signal peptide (SP) prediction across all domains of life and distinguishes between three types of prokaryotic SPs, i.e., SP (Sec/SPI): standard secretory signal peptides transported by the Sec translocon and cleaved by Signal Peptidase I, Sec/SPII (LIPO): lipoprotein signal peptides transported by the Sec translocon and cleaved by Signal Peptidase II and Tat/SPI (twin-arginine translocation pathway, TAT): signal peptides transported by the Tat translocon and cleaved by Signal Peptidase I. A total of 392 proteins were predicted to have one of these types of signal peptide in *M. tuberculosis* proteome (207 SP, 113 LIPO and 72 TAT). Of those we identified 140 in CFP (62 SP, 53 LIPO and 25 TAT), being many of them well recognized secreted proteins, particularly FbpA (Rv3804c), FbpB (Rv1886c), FbpC (Rv0129c), Apa (Rv1860), Mpt64 (Rv1980c), PstS1 (Rv0934), LpqH (Rv3736), among others (S2 Table).

This approach allowed for the identification of proteins targeted to the signal-sequence-dependent secretory pathways. To export proteins across its unique cell wall, mycobacteria utilize the general secretion pathways, twin-arginine transporter, and up to five distinct ESX secretion systems (designated ESX-1 through ESX-5, referred to as the type VII secretion system: T7SS), which various functions in virulence, iron acquisition, and cell surface decoration [14]. The ESX-1 system was the first of the T7SS to be identified and is responsible for the secretion of EsxA (6 kDa early secretory antigenic target, ESAT-6, Rv3875) and EsxB (Rv3874) [47]. It is important to note that proteins belonging to ESX secretion systems gene clusters as well as closely related PE and PPE gene families are *M. tuberculosis* secreted proteins that do not have classical secretion signals [15,48]. Taken this into consideration, we identified in CFP several proteins of ESAT-6 family: EsxA (Rv3875), EsxB (Rv3874), EsxG (Rv0287), EsxI (Rv1037c), EsxK (Rv1197) grouped with EsxP (Rv2347c) and EsxJ (Rv1038c), EsxL (Rv1198), EsxN (Rv1793) grouped with EsxV (Rv3619c), EsxO (Rv2346c) and EsxW (Rv3620c). None of those were predicted by SignalP to contain a signal peptide. Besides, various proteins of ESX-1 secretion system detected in this analysis were not predicted to have a signal peptide, including EspA (Rv3616c), EspD (Rv3614c), EspC (Rv3615c) and EspB (Rv3881c). All of them count with experimental evidence of being secreted [32]. Finally, we detected 8 PE and PPE family proteins in our sample, from which 3 were predicted to have a signal peptide, i.e., PE13 (Rv1195), PE5 (Rv0285) and PE15 (Rv1386) and 5 were not predicted to have a signal peptide, i.e., PE25 (Rv2431c), PE31 (Rv3477), PPE41 (Rv2430c), PPE18 (Rv1196) and PPE60 (Rv3478). In particular, PE25 and PPE41 form a heterodimer that is secreted by the ESX-5 system of *M. tuberculosis* [49]. In summary, various proteins with signal peptides were detected in our sample and several other proteins related to T7SS were identified. The SignalP 5.0 server was a suitable approach in order to predict secreted proteins with classical signal peptides but it has limitations to analyze proteins bearing non-classical secretion signals.

### Integrative analysis with previous proteomic studies

In order to get more information on the results obtained and validate them, former research studies, which used different and complementary approaches to characterize *M. tuberculosis H37Rv* CFP, were compared against our results. We selected relevant previous proteomic studies reporting a similar methodology of mycobacterial culture and CFP preparation [7,13,37]. Malen *et al.* characterized a culture filtrate of *M. tuberculosis* H37Rv, considerably enriched for secreted proteins, with two complementary approaches (i) 2D gel electrophoresis combined with MALDI-TOF MS and (ii) LC coupled MS/MS. Peptides derived from a total of 257 proteins were identified, of which 254 were annotated with an Rv identifier [7]. Later, de Souza *et al.* using nano-LC in tandem with an Orbitrap mass spectrometer performed a proteomic screening to identify proteins in culture filtrate, membrane fraction and whole cell lysate of *Mycobacterium tuberculosis*. Through this approach they identified 2182 different proteins in the different fractions, specifically 458 proteins in CFP, 1447 in the membrane fraction and 1880 in the whole cell lysate [13]. In a recent report, Albrethsen *et al*. used label-free LC-MS/MS of SDS-PAGE fractionated samples to investigate the culture filtrate proteome of *M. tuberculosis* H37Rv bacteria in normal log-phase growth and after 6 weeks of nutrient starvation. In total, in this study 1362 proteins were identified in six CFP samples analyzed (three log phase samples and three 6-week-starved CFP samples) [37]. The comparison of proteins identified in our analysis against the proteins identified in CFP of these former proteomic researches showed a common group of 122 proteins consistently detected (Fig 3A). Among these proteins, 41 belong to the P90% subgroup indicating that these are highly abundant proteins. The most important proteins of this common group include 10 kDa chaperonin GroES (Rv3418c), ESAT-6-like protein EsxB (Rv3874), 6 kDa early secretory antigenic EsxA (Rv3875), Chaperone protein DnaK (Rv0350), the secreted antigen 85 complex −85A (Rv3804c), 85B (Rv1886c) and 85C (Rv0129c)-, Glutamine synthetase GlnA1 (Rv2220), Immunogenic protein Mpt64 (Rv1980c), Superoxide dismutase SodA (Rv3846), Thioredoxin TrxA (Rv3914), Glycogen accumulation regulator GarA (Rv1827), Phosphate-binding protein PstS1 (Rv0934), Alanine and proline-rich secreted protein Apa (Rv1860) and various other ESAT-6 family proteins (EsxO Rv2346c, EsxL Rv1198, EsxG Rv0287). Moreover, 1073 proteins were shared between our set of proteins and the list reported by Albrethsen *et al.* [37], representing 81.7% of the proteins identified by us and confirming a strong concordance between both analysis.

**Fig 3.**
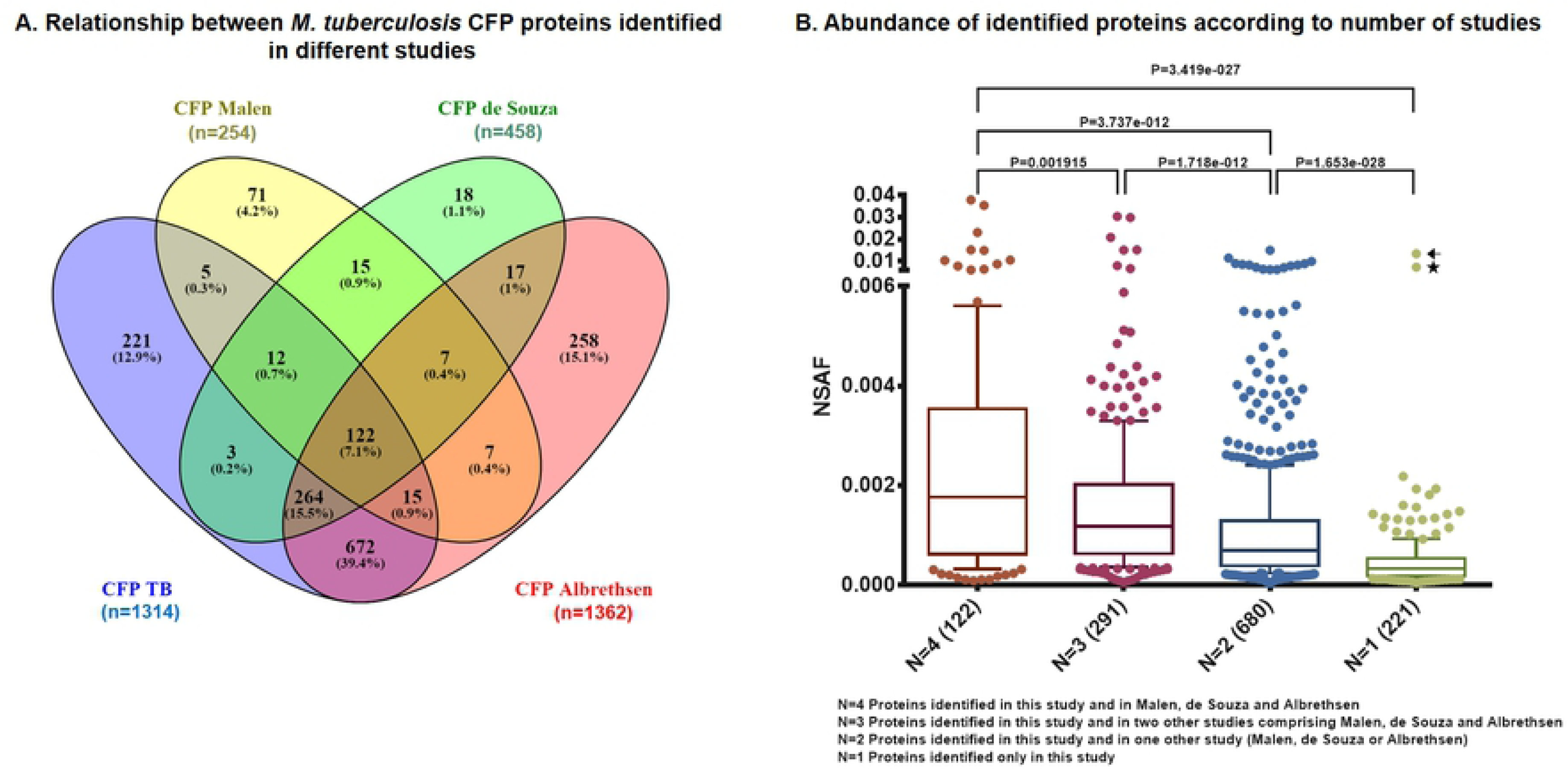
Comparison of *M. tuberculosis* CFP with other relevant proteomic studies. (A) Analysis of *M. tuberculosis* CFP protein list (CFP TB: this study) versus other relevant proteomic studies of *M. tuberculosis* CPF, identified as CPF Malen [7], CFP de Souza [13] and CFP Albrethsen [37] by Venn Diagram comparison (Venny’s on-line reference [38]). (B) Protein abundance estimation of proteins identified this study (CFP TB) and in all of the three other studies evaluated (N=4), in this study and in two other studies (N=3), in this study and in one other study (N=2), or only in this study (N=1). The arrow indicates the protein Rv3620c (esxW) that was identified in an additional study [8] not included in the comparison of Fig 3A. The star indicates the protein Rv3118 (sseC1) which has an identical (100% identity) second copy at Rv0814c (sseC2) which was identified in CFP Albrethsen [37]. p-value obtained after Mann-Whitney test comparison of the median of two groups is shown, and the groups compared in each case is indicated with a line above each graph.

The label free quantitative approach applied in this study was exploited to compare the abundance in our sample of proteins identified in all the studies included in the analysis (N=4) versus those proteins identified in 3 (N=3), 2 (N=2) or 1 study (only this study) (N=1). Fig 3B clearly shows that proteins identified in the four studies are on average more abundant than proteins identified in the other groups analyzed. Moreover, proteins identified in at least 2 studies (N=3 or N=2) are globally more abundant than proteins identified exclusively in the present work.

Additional analysis comparing our data against the proteomic quantitative approach performed by de Souza *et al* [13] allowed us to identify a subgroups of highly represented proteins consisting of those identified in this work and also in the three fractions studied by this previous work, i.e. culture filtrate, membrane fraction and whole cell lysate. This subgroup accounted for 43.2% of protein abundance expressed as NSAF in this work and 29.2% of emPAI calculated by the cited research. Besides, a group of 921 proteins identified in membrane fraction and/or whole cell lysate prepared by de Souza *et al* and accounting for 13.3 % of calculated emPAI was not detected in the culture filtrate prepared by them neither in CFP prepared in this study [13]. These results are summarized in S3 Table.

As a whole these results show that the CFP prepared in the present work exhibited a good correlation with previous studies, both in terms of qualitative proteomic composition as well as in relation to the quantitative estimation of protein abundance. Proteins highly represented in our sample are proteins either frequently identified by others using complementary approaches in culture filtrates of MTB, and thus confirming that our sample is enriched in proteins that the bacteria does secrete, or ubiquitously detected in different *M. tuberculosis* cellular fractions, indicating that these could represent highly expressed proteins.

Finally, with this approach 30 proteins not previously annotated with proteomic data in Mycobrowser website (Release 3 (2018-06-05)) [33] were identified (S4 Table). This list, principally composed by proteins classified as conserved hypotheticals, includes the ESX-3 secretion-associated protein EspG3 (Rv0289) identified with 4 unique peptides in CFP(1) and 5 unique peptides in CFP(2) and the Two component sensor histidine kinase DosT (Rv2027c) identified with 2 unique peptides in each replicate. Further comparison of these proteins with the results obtained in a proteome-wide scale approach based on SWATH mass spectrometry [50] allow us the identification, to the best of our knowledge, of 8 proteins without previous evidence of expression at the protein level. In S5 Table these proteins are listed as well as the scans of their corresponding peptides.

### O-glycosylation analysis

To complement our analysis, the presence of the most common naturally occurring glycan residues in mycobacteria was analyzed: hexoses, like mannose, glucose or galactose, which are highly reported in mycobacterial lipoproteins [18], deoxyhexoses, like fucose and rhamnose, that are important components of the cell surface glycans [51], the pentose sugar arabinose also reported in some glycoproteins [18] and as part of the mycolyl-arabinogalactan-peptidoglycan of the cell wall [52], and heptoses, recognized to be transferred by heptosyltransferases using ADP-heptose [53]. Our rationale was that the nano LC MS/MS technology used in this work, by having more than four orders of magnitude intrascan dynamic range and a femtogram-level sensitivity, would allow the direct identification of modified peptides, without previous affinity-based strategies for glycosylated protein enrichment.

In each replica several O-glycosylation events were detected and after comparing them a reduced subgroup of common peptides and proteins was defined and selected for further analysis. O-glycosylation profile analysis revealed the presence of 154 common glycosylation events in 135 common modified peptides in both replicas of MTB culture filtrate (Table 2). The O-glycosylated common peptides were identified in 363 scans, consisting in at least 2 scans per peptide (1 scan per replica) and a maximum of 8 scans in the case of Hex-Hex-Hex modification of Alanine and proline rich secreted protein Apa (Rv1860) (S6 Table). The four studied monosaccharide modifications (Hex, Pentose, DeoxyHex and Heptose) were highly similarly represented in culture filtrate proteins, being Hex the most frequent modification (Table 2). In many cases the unmodified peptide was identified along with the modified peptide, indicating that glycosylated and unglycosylated proteins isoforms are present (S2 Fig), as was previously reported for the conserved lipoprotein LprG [54].

**Table 2.**
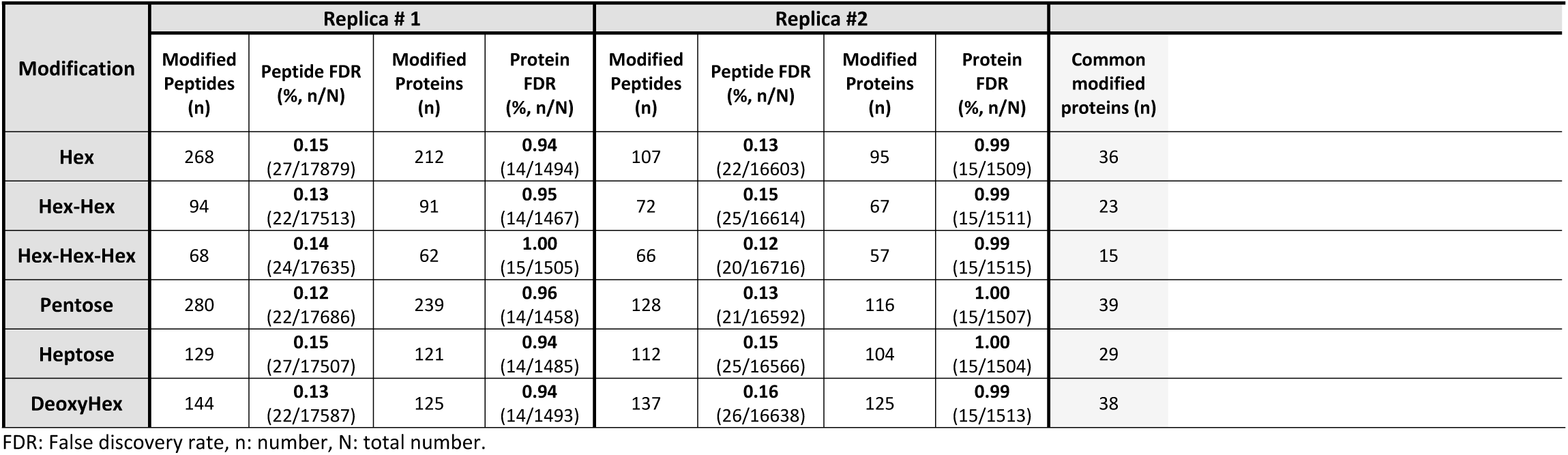
O-glycosylation profile of M. tuberculosis culture filtrate proteins identified by LC MS/MS.

| Modification | Replica # 1 |  |  |  | Replica #2 |  |  |  | Common modified protein |
| --- | --- | --- | --- | --- | --- | --- | --- | --- | --- |
|  | Modified Peptides (n) | Peptide FDR (% , n/N) | Modified Proteins (n) | Protein FDR (% , n/N) | Modified Peptides (n) | Peptide FDR (% , n/N) | Modified Proteins (n) | Protein FDR (% , n/N) |  |
| **Hex** | 268 | **0.15**<br>(27/17879) | 212 | **0.94**<br>(14/1494) | 107 | **0.13**<br>(22/16603) | 95 | **0.99**<br>(15/1509) | 36 |
| **Hex-Hex** | 94 | **0.13**<br>(22/17513) | 91 | **0.95**<br>(14/1467) | 72 | **0.15**<br>(25/16614) | 67 | **0.99**<br>(15/1511) | 23 |
| **Hex-Hex-Hex** | 68 | **0.14**<br>(24/17635) | 62 | **1.00**<br>(15/1505) | 66 | **0.12**<br>(20/16716) | 57 | **0.99**<br>(15/1515) | 15 |
| **Pentose** | 280 | **0.12**<br>(22/17686) | 239 | **0.96**<br>(14/1458) | 128 | **0.13**<br>(21/16592) | 116 | **1.00**<br>(15/1507) | 39 |
| **Heptose** | 129 | **0.15**<br>(27/17507) | 121 | **0.94**<br>(14/1485) | 112 | **0.15**<br>(25/16566) | 104 | **1.00**<br>(15/1504) | 29 |
| **DeoxyHex** | 144 | **0.13**<br>(22/17587) | 125 | **0.94**<br>(14/1493) | 137 | **0.16**<br>(26/16638) | 125 | **0.99**<br>(15/1513) | 38 |

FDR: False discovery rate, n: number, N: total number.

O-glycosylation modification were detected in 108 different MTB culture filtrate proteins, 52 of them presented at least 3 scans of the modified peptide and 12 bore more than one of the searched modifications, i.e. Apa (Rv1860), EsxA (Rv3875), LpqH (Rv3763), LppO (Rv2290), CarB (Rv1384), AceE (Rv2241), FhaA (Rv0020c), PstS1 (Rv0934), LprF (Rv1368), DsbF (Rv1677), Mpt64 (Rv1980c) and DevR (Rv3133c) (Fig 4A). What is Interesting to highlight is the high number of scans of modified peptides corresponding to Apa (Rv1860), most of them corresponding to Hex, Hex-Hex or Hex-Hex-Hex. This protein, also known as immunogenic protein MPT32 or 45-kDa glycoprotein is a largely characterized secreted mannosylated glycoprotein [55] and in agreement with previous reports we found scans corresponding to the presence of one, two or three hexoses between T313, T315, T316 and T318 as glycosylation sites [21]. It is currently believed that mannosylated proteins can act as potential adhesins and it was demonstrated that Apa is associated with the cell wall and binds lung surfactant protein A (SP-A) and other immune system C-TLs containing homologous functional domains [56]. The 19 kDa lipoprotein antigen precursor LpqH (Rv3763), also showing an important number of Hex-Hex and Hex-Hex-Hex modified peptides, is a well-known glycosylated protein exposed in the bacterial cell envelope, that was postulated to be used by mycobacteria to enable their entry into the macrophage through interaction with mannose receptors (MRs) of this host cells [57].

**Fig 4.**
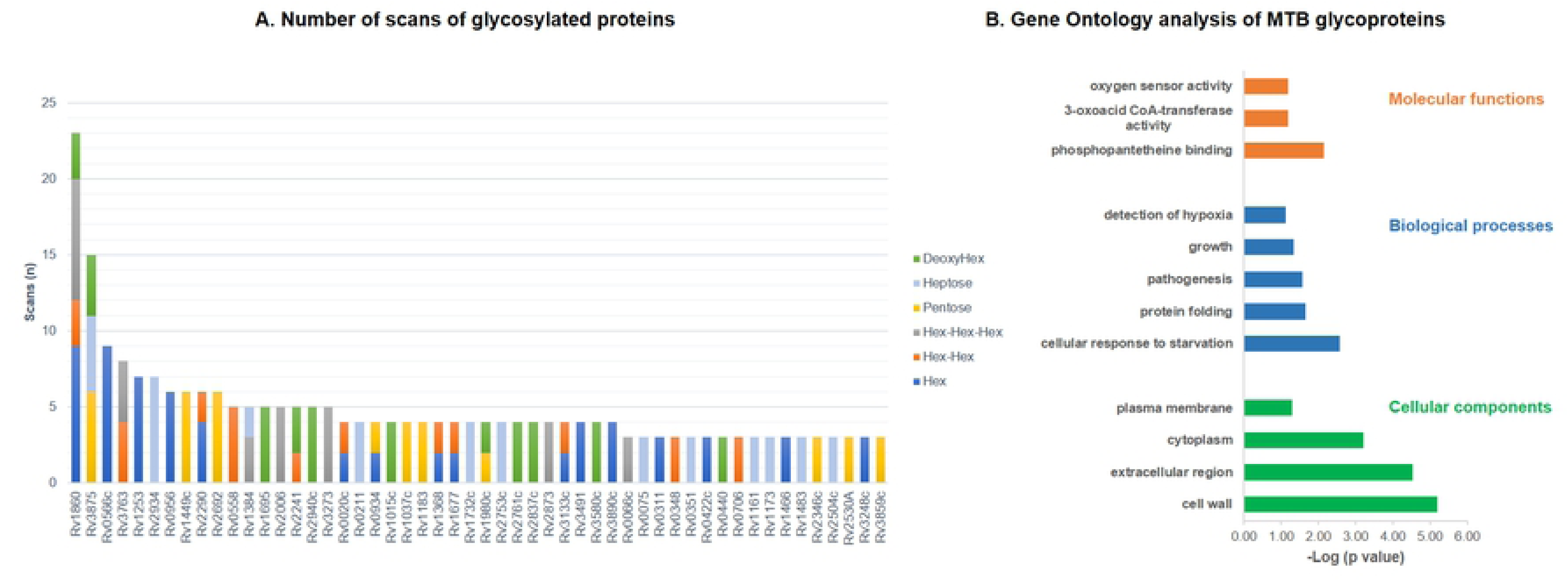
Description of O-glycosylated proteins in *M. tuberculosis* CFP. (A) Scans of O-glycosylated peptides identified in MTB culture filtrate proteins. Each analyzed modification is displayed with a different bar color. Individual scans of both replicates were considered and only 52 proteins identified by at least three different scans are shown in the graph. (B) Gene Ontology analysis of MTB culture filtrate glycoproteins. Principal categories of enriched terms (p<0.05) obtained analyzing proteins with common glycosylation in both replicates with David Gene Functional Classification Tool [39,40] using Molecular Functions, Biological Processes and Cellular Component Ontology database and *M. tuberculosis* H37Rv total proteins as background.

It is important to note that the precise O-glycosylation site assignation is hampered by the fact that collision energies used for peptide fragmentation cause the breakage of the weaker O-glycosydic bond leaving behind mostly unmodified fragments. Although the glycosylation site assignation was not the aim of our study, the utility XDScoring of Patternlab for proteomics developed for statistical phosphopeptide site localization [58], was preliminary tested in our data. Glycosylation site p-value is presented in S6 Table.

Glycosylation plays a significant role in MTB adaptive processes and in particular cell-cell recognition between the pathogen and its host is mediated in part by glycosylated proteins. Based on the Gene Ontology (GO) analysis of the glycoproteins identified, cellular response to starvation, protein folding and pathogenesis were highly enriched biological processes. Our GO analysis further showed that most of the glycoproteins identified were localized in the cell wall and extracellular region and that phosphopantetheine binding (including Mas, Pks2, PpsD, Pks13 and Pks5), 3-oxoacid CoA-transferase activity and oxygen sensor activity (DesV and DesR) were significantly enriched molecular function categories (Fig 4B).

### O-glycosylation validation

Of the 108 identified glycoproteins 21 were identified as candidate glycoproteins in the lectin interacting enriched membrane protein using WGA-affinity capture [12]. Besides, 12 glycoproteins bearing mono- or polyhexose modifications in our analysis have been included in a recent review of protein glycosylation and lipoglycosylation in *M. tuberculosis* [18], where experimental evidence was summarized. Among them several lipoproteins are included: LprA (Rv1270c), LprF (Rv1368), LppO (Rv2290), LpqH (Rv3763), PstS1 (Rv0934) and Mpt83 (Rv2873). Moreover, four of these proteins were consistently found with the same type of hexose O-glycosylation in culture filtrate of MTB, i.e. Apa (Rv1860), LppO (Rv2290), Rv2799 and Rv3491 [21]. Our results confirm the presence of glycosylated lipoproteins in culture filtrate aiding to the growing evidence for glycosylation of mycobacterial lipoproteins [18,21]. Besides, we identified mono- or polyhexose modifications in DsbF (Rv1677), a probable conserved lipoprotein. The same DsbF glycosylation pattern was reported in a recent glycoproteomic analysis of MTB cell lysates of four different linages [17]. In this work 27 proteins of our list were also described as O-glycosylated, including HtrA (Rv1223), Wag31 (Rv2145c), FbpB (Rv1886c) and Rv2411c.

To further evaluate the reproducibility of our results and validate them we looked for O-glycosylated proteins in the raw data files deposited by Albrethsen *et al*. [37] at the ProteomeXchange Consortium. By means of this approach 22 proteins with the same O-glycosylation type were found and after peptide sequence comparison we confirmed 20 modified peptides in common with our results, corresponding to 11 different proteins LprA (Rv1270c), DsbF (Rv1677), Rv1732c, Apa (Rv1860), AroE (Rv2552c), Rv2799, Mpt83 (Rv2873), SahH (Rv3248c), Rv3491, LpqH (Rv3763) and EsxA (Rv3875). The scans corresponding to these peptides are presented in S7 Table.

As a whole, we are reporting 62 novel O-glycosylated proteins including hexose, heptose, pentose or deoxyhexose, 10 of them being validated with raw data re-analysis of the selected previous work [37]. Several relevant scans corresponding to glycosylated peptides were statistically confirmed in Mascot Server MS/MS Ions Search against NCBIprot (AA) database of all taxonomies [41] (S3 Fig). Interestingly EsxA (Rv3875) was found with three different types of O-glycosylation - DeoxyHex, Pentose and Heptose – (Fig 4A). Of those the presence of two heptoses, one in T61 and the other in T63 was also identified in at least one replica of log phase culture filtrates in Albrethsen *et al*. [37] (S7 Table). A representative peptide spectrum of this modification including peptide ions fragment matches is shown in Fig 5. EsxA (Rv3875) and its chaperone protein EsxB (Rv3874), localized in Region of Difference 1 (RD1) of the MTB genome, are important virulence factors of MTB and the most immunodominant antigens thus far identified [59]. EsxA (or ESAT-6) is included in several vaccine candidates in development [60] and is also the core antigen in the IFN-γ release assays (IGRA) used to diagnose latent infection [61]. A former report described that an N-terminal Thr acetylation (+42Da) was identified in some species of this protein obtained in a short-term MTB culture filtrate [62] and other literature mentioned this protein as being glycosylated [17,63], however, to our knowledge, we are presenting novel evidence of several O-glycosylation events in this relevant secreted antigen.

**Fig 5.**
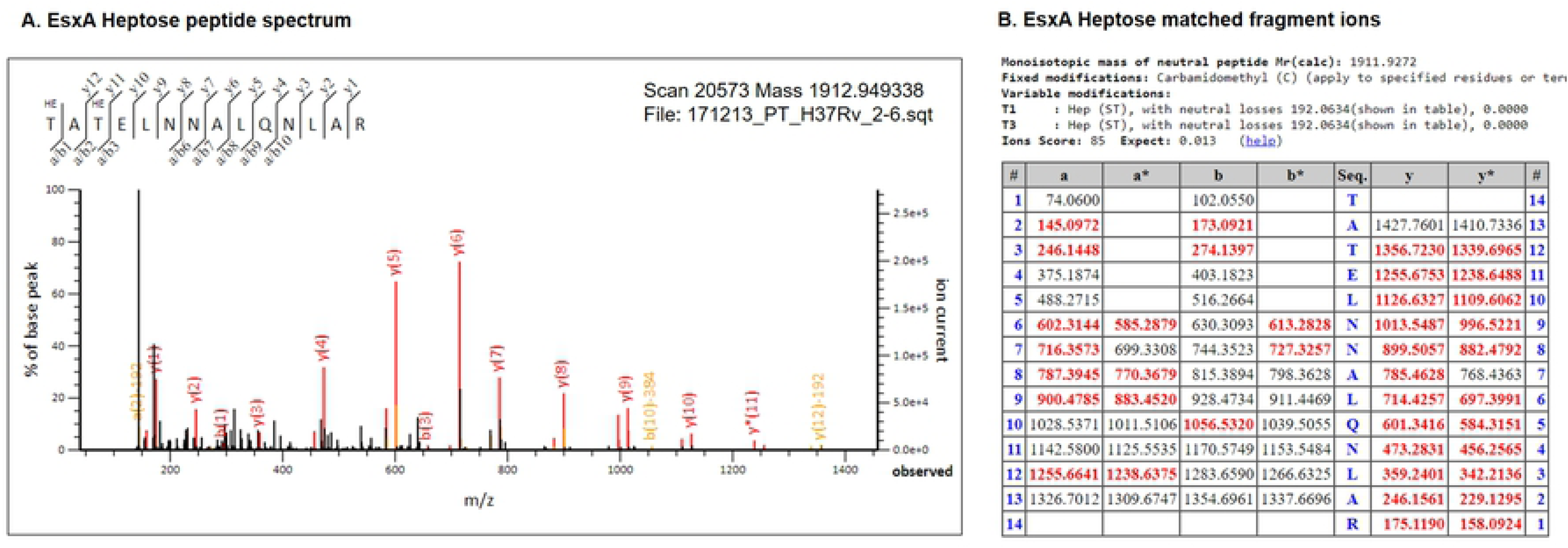
EsxA heptose-modified peptide spectra. (A) Representative spectrum of EsxA heptose-modified peptide statistically confirmed by Mascot Server MS/MS Ions Search (HE = heptose). (B) Fragment ions matches indicated in bold red as reported in Mascot Server.

## Conclusion

Membrane and exported proteins are crucial players for maintenance and survival of bacterial organisms in infected hosts, and their contribution to pathogenesis and immunological responses make these proteins relevant targets for medical research [11]. Consistently, various of the proteins identified in *M. tuberculosis* CFP were proposed as relevant mycobacterial virulence factors [64], putative active infection biomarkers [9] or vaccine candidates [60,65]. This shotgun proteomic approach allowed a deep comprehension of *M. tuberculosis* H37Rv culture filtrate proteins reporting proteomic evidence in this sub-fraction for 1314 proteins. In that sense it is important to note that although this method is highly sensitive, specificity was prioritized by selecting as post-processing criteria that considered only proteins with at least two different peptide spectrum matches.

In addition to proteins that have not been previously reported in *M. tuberculosis* H37Rv CFP, we also found proteins consistently detected in previous proteomic studies which were further confirmed as highly abundant proteins. Many of these proteins were previously described in culture filtrates of MTB or detected in different *M. tuberculosis* cellular fractions, including membrane fraction and whole cell lysate. This could suggest that two complementary pathways are accounting for our observations. On one hand, the abundance of certain proteins in CFP appear to be truly related to protein-specific export mechanisms, while on the other hand the occurrence of some proteins in CFP due to bacterial autolysis in combination with high levels of protein expression and extracellular stability cannot be ruled out. Nevertheless, the GO ontology Cellular Component analysis and the integrative analysis performed with relevant research papers confirms that our sample is indeed enriched in proteins that the bacteria secretes to the extracellular space.

Supporting this, we could identify several proteins with predicted N-terminal signal peptide indicating that these are targeted to the secretory pathways [66], as well as various proteins belonging to the ESX secretion systems, and PE and PPE families known to be secreted by T7SS, but recognized as not to have classical secretion signals [48].

With the aim to assess the role of protein O-glycosylation in MTB virulence and host-pathogen interactions [16,18], this study described the identification of 154 glycosylation events in 108 MTB proteins. In particular, several lipoproteins were found glycosylated in culture filtrate. Lipoproteins have been shown to play key roles in adhesion to host cells, modulation of inflammatory processes, and translocation of virulence factors into host cells [67]. The growing evidence of glycosylation of mycobacterial lipoproteins including the results presented here, indicates that glycosylation plays a significant role in the function and regulation of this group of proteins. Along with lipoproteins, other relevant glycoproteins identified were mainly involved in cellular response to starvation, protein folding and pathogenesis. As a novel contribution of this work, we are reporting that the virulence factor EsxA is glycosylated in MTB culture filtrate. It is important to note that in addition to EsxA other glycosylated proteins identified in this work have been proposed as diagnostic biomarkers for TB active disease. Protein glycosylation data presented here, including the coexistence of related protein glycoforms evidenced in this work, should be considered for designing antibody-based diagnostic test targeting *M. tuberculosis* antigens. Besides, as reported for other pathogens [68,69], protein glycosylation diversity could be a key mechanism to provide antigenic variability aiding in the immune subversion of this pathogen.

Our study provided an integrative evaluation of MTB culture filtrate proteins, bringing evidence of the expression of some proteins not previously detected at protein level, and confirming and enlarging the database of O-glycosylated proteins. This novel information may raise new questions on the role of protein O-glycosylation on the biology of MTB, as well as it will contribute to complement the knowledge of its relevant biomarkers, virulence factors and vaccine candidates.

## Acknowledgments

We thank Rosario Duran (IIBCE/Institut Pasteur de Montevideo, Uruguay) for critical reading of the manuscript and helpful discussion about LC MS/MS experiment design and data analysis. We also thank Alejandro Leyva (Institut Pasteur de Montevideo, Uruguay) for technical assistance with Orbitrap mass spectrometer and Paulo C. Carvalho (Fiocruz, Brazil) for his valuable collaboration with data analysis in PatternLab for Proteomics. Finally, we would like to thank the staff of Comisión Honoraria de Lucha Antituberculosa y Enfermedades Prevalentes (Montevideo, Uruguay), for technical assistance with *M. tuberculosis* culture.

## Supporting information

**S1 Fig. Analysis of *M. tuberculosis* CFP by liquid chromatography tandem mass spectrometry (LC-MS/MS).**

**2A**: *M. tuberculosis* CFP analysis by 1D SDS-PAGE 15% and CCB G-250 staining. Two technical replicates (CFP(1) and CFP(2), 25 ug each) were loaded. Six gel slices were excised from each lane according to protein density. Numbers indicate gel slices analyzed by LC-MS/MS. MWM: Molecular weight marker (Thermo Fischer Scientific, # 26616). **2B:** Spectrum counts of proteins identified in each technical replicate. Replicates show no statistical differences (p>0.05). **2C:** Analysis of proteins identified in each replicate by area-proportional Venn Diagram comparison [31]

**S2 Fig. Proteins showing glycosylated and unglycosylated equivalent peptides.**

Some protein examples are shown: 1) Apa (modification: Hex), 2) LprF (modification: Hex), LppO (modification: Hex-Hex), Apa (modification: Hex-Hex-Hex), EsxA (modification: Pentose).

**S3 Fig. Scans of glycosylated peptides statistically confirmed in Mascot Server MS/MS Ions Search against NCBIprot (AA).**

Some examples are shown: 1) LppO (modification: Hex), 2) EsxA (modification: DeoxyHex), 3) EsxA (modification: Pentose).

**S1 Table. Proteins identified with nano-HPLC MS/MS.**

Sheet 1) Common proteins list including Uniprot identification, protein description, protein length and molecular weight, gene name and *M. tuberculosis H37Rv* gene annotation (Rv) of Sanger Institut (http://sanger.ac.uk/projects/M_tuberculosis/Gene_list/). Sheet 2) Proteins identified in replica CFP(1), Sheet 3) Proteins identified in replica CFP(2), both lists including Uniprot identification as obtained in Patternlab for Proteomics, sequence count, spectrum count, number of unique peptides, protein coverage and protein description.

**S2 Table. Proteins with predicted signal peptides**

Sheet 1) Signal peptide prediction (SignalP 5.0) in *M. tuberculosis* H37Rv reference proteome (UP000001584), Sheet 2) Signal peptide prediction (SignalP 5.0) in *M. tuberculosis* H37Rv CFP, Sheet 3) Proteins in *M. tuberculosis* H37Rv CFP with signal peptides predicted with SignalP 5.0.

**S3 Table. Protein abundance comparison against de Souza *et al*, 2011**

Comparison of our proteomic data against the proteomic quantitative approach performed by de Souza *et al*, 2011 [13].

**S4 Table. Proteins without proteomic annotation in Mycobrowser**

Proteins identified in *M. tuberculosis* H37Rv CFP without proteomic annotation in Mycobrowser (Release 3 (2018-06-05)) [33].

**S5 Table. Peptides of proteins not previously detected at proteomic level**

Sheet 1) Proteins in *M. tuberculosis* H37Rv CFP without previous evidence of expression at protein level, Sheet 2) Scans of peptides confirming proteins identified in *M. tuberculosis* H37Rv CFP without previous evidence at protein level.

**S6 Table. Scans of O-glycosylated peptides in *M. tuberculosis* H37Rv culture filtrate proteins**

The table includes the File name where the scan was identified, the scan number, peptide charge (Z), measured and theorical mass and the difference (in ppm), scores (primary, secondary, etc), peptide sequence, modification (glycan), glycosylation site p-value, protein and gene data.

**S7 Table. O-glycosylation analysis of raw files of Alberthsen *et al*, 2013.**

Scans confirming O-glycosylated peptides identified by us in the analysis of the raw data files deposited by Albrethsen *et al*. [37].

